# A locomotion-based assay to measure prepulse inhibition in zebrafish larvae

**DOI:** 10.1101/2023.09.29.560094

**Authors:** Emily Read, Robert Hindges

## Abstract

Sensory gating, assessed using a prepulse inhibition assay (PPI), is a promising endophenotype of neuropsychiatric disorders that can be measured in larval zebrafish models. However, current PPI assays require high-speed cameras to capture rapid c-bend startle behaviours. In this study, we designed and employed a PPI paradigm that uses locomotion as read-out of startle responses. PPI percentage was measured at a maximum of 86% and this was reduced to 42% upon administration of an NMDA receptor antagonist, MK-801. This work provides the foundation for simpler and more accessible PPI assays using larval zebrafish to model key endophenotypes of neurodevelopmental disorders.

## Introduction and Results

In the field of neuropsychiatric disorders, impairment of startle response habituation and sensitisation has been proposed as one of the most promising translational endophenotypes for various disorders, including schizophrenia. Measurement of this endophenotype is robust via the prepulse inhibition (PPI) assay. Here, startle responses are reduced in trials when a lower intensity (prepulse) stimulus is presented 30-500 msec prior to a higher intensity (startle) stimulus. This is a measure of sensory gating: the ability of the autonomic nervous system to filter sensory information. Patients with schizophrenia and high-risk individuals show reliable PPI reductions compared to healthy controls, maintaining a high startle response even if the startle stimulus is preceded by a prepulse (Li et al., 2021; San-Martin et al., 2020; Swerdlow et al., 2006; Takahashi & Kamio, 2018; Ziermans et al., 2012). PPI is detected across different species, including common model organisms, and therefore the assay has become highly relevant to investigate mechanisms underlying the aetiology of disorders.

PPI assays can be modelled in the zebrafish (*Danio rerio*). Using larval stages, the proportion of fish displaying a characteristic ‘c-bend’ startle response (a very fast escape/startle reaction) is reduced when the startle stimulus is preceded by a prepulse (Burgess & Granato, 2007). The parameters required for PPI in zebrafish larvae are strikingly similar to that used in human PPI assays (Wolman et al., 2011). Furthermore, pharmacological and genetic manipulations, that are used to model schizophrenia in the zebrafish, also lead to reductions in the PPI response (Bergeron et al., 2015; Burgess & Granato, 2007), as is observed in schizophrenia.

However, the c-bend startle must be measured using high-speed cameras, as it occurs in just 20 msec or less and requires detection of precise angular changes in the orientation of the larva’s tail with respect to its head (Bergeron et al., 2015). Due to the high expense of such equipment and the more extensive data analysis, the PPI assay can be difficult to implement. More recent studies have used other behavioural read-outs of startle responses, including locomotion which can be done at camera frame rates of just 30fps, in the context of habituation in larvae, or general motion detection in adult fish for sensory gating experiments (Beppi et al., 2021; Kirshenbaum et al., 2019). However, such simpler approaches have not yet been created for a PPI paradigm in larvae or in context with schizophrenia-relevant pharmacological modulation. In this report, we present the establishment of a PPI assay based on only measuring locomotion startle responses in zebrafish larvae and furthermore show that it is possible to observe a schizophrenia-reminiscent PPI reduction by administration of the glutamatergic NMDA receptor antagonist, MK-801.

The locomotion-based PPI assay was set up using the Zantiks MWP (https://zantiks.com/), which is an automated system containing a low-speed camera at 30fps and 720 x 540 pixel resolution. To produce the startle stimuli, vibrations were delivered using an built-in motor which is script-controlled. To generate a reliable PPI protocol using this system, we optimised several parameters, namely vibration intensity and time intervals between stimulus presentations.

Based on previous literature, vibrations eliciting a variation in startle responses in zebrafish larvae are between 100-1000Hz (Beppi et al., 2021; Bergeron et al., 2015; Burgess & Granato, 2007). As such, we set the frequency of both the startle and prepulse stimuli to 200Hz. To deliver the vibration, the motor was set to move by either 1.8°(1 full step), 0.9° (1/2 step), 0.45° (1/4 step), or 0.225°(1/8 step) in 4 x 5 msec clockwise and anti-clockwise movements. To evaluate if larvae would respond differently to different motor step sizes, four single pulses (one at each step size), spaced at 5-minute inter-trial intervals (ITIs) were presented to larvae. Previous studies have used a minimum of 15 seconds (Burgess & Granato, 2007) and a maximum of 15 minutes (Beppi et al., 2021) ITI, thus we felt confident that no habituation to startle stimuli would be observed at 5 minutes. Startle response magnitude and likelihood were measured using distance moved (locomotion) per 100 msecs.

Larvae showed a graded startle likelihood at varying pulse intensities (Figure 1A). 200Hz pulses at 1 step and ½ step showed a response likelihood of 0.82, at ¼ step it was 0.80, and at 1/8 step this decreased to 0.52. Analysing the response magnitudes with a one-way ANOVA showed a similar pattern, that the total distance larvae moved in response to the startle was different across pulse intensities (*F*(3, 279) = 10.67, *p* < 0.0001; Figure 1B). There was a significantly greater startle magnitude to 1 step compared to ½ step (*p* < 0.0001), ¼ step (*p* = 0.0196) and 1/8 step (*p* = 0.0003). PPI in the larval zebrafish has previously been defined as a difference in response *probability* when the startle stimulus is preceded by a prepulse stimulus (Burgess & Granato, 2007). To generate a consistent decrease in response probability, the startle vibration was set at 200Hz with one full motor step, and the prepulse vibration set to 200Hz with 1/8 motor step. These startle and prepulse vibrations were applied to all subsequent PPI assays.

**Figure 1.**
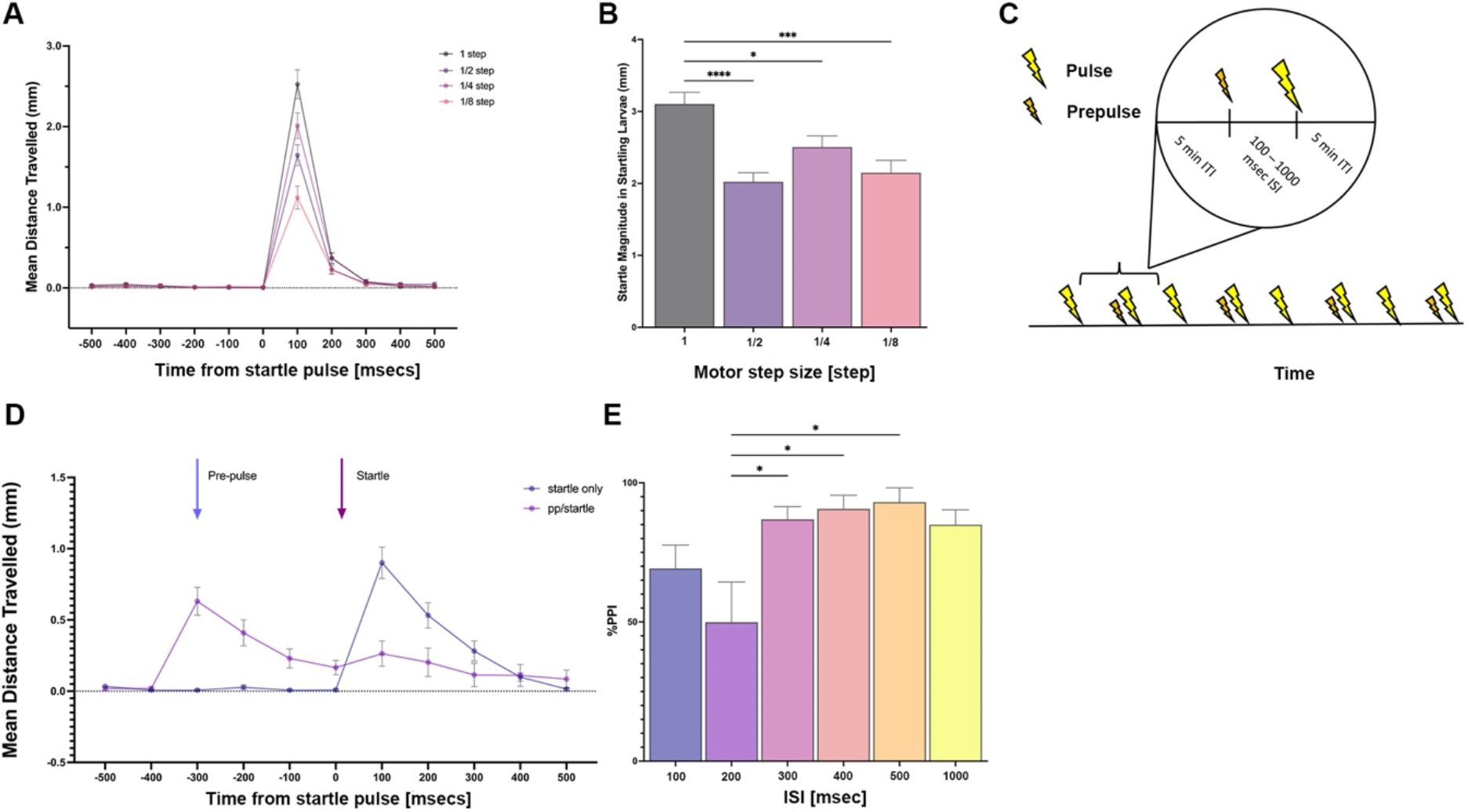
Locomo.on-based prepulse inhibi.on. **A**. The startle response of zebrafish larvae (6dpf) after within-subject presentation of four single 200Hz 20msec vibrations (one vibration per motor step magnitude, randomised order), presented as average locomotion across larvae for each pulse intensity. **B**. Startle magnitude of responder larvae, measured as average locomotion across larvae for each pulse intensity. **C**. Diagram to show the finalised PPI assay procedure. **D**. 300 msec ISI visualised locomotion data: mean distance travelled by larvae in startle pulse only and prepulse/startle trials. Arrows indicate time points where the prepulse and startle stimuli are delivered on relevant trials. **E**. Percentage PPI of responder larvae at inter-stimulus intervals (ISIs) of 100 – 1000 msec. For all *N*-numbers, see Table 1 in Methods section. *\** = *p*<0.05, *** = *p*<0.001, **** = *p*<0.0001.

When running the PPI assay in full, we used the above parameters but varied the inter-stimulus interval (ISI) between prepulse and startle stimuli between 100 to 1000 milliseconds (Figure 1C). This is consistent with earlier findings showing that the ISI greatly impacts PPI percentage between 30 to 3000msec, with the highest PPI percentages being observed at intermediate ISIs around 300-500msecs (Bergeron et al., 2015; Burgess & Granato, 2007). In total, eight vibration trials were delivered in each PPI assay. Startle pulse alone and prepulse/startle trials were evenly interspersed with a total of four of each trial type (Figure 1C).

**Table 1.** *N*-numbers per group in each vibration/PPI experiment.

|  | ***N*-total per group**<br>**(after exclusions)** | ***N*- responder larvae**<br>**per group (startle)** | ***N*- responder larvae**<br>**per group (pp/startle)** |
| --- | --- | --- | --- |
| **Single Pulses (Figure 1A/B)** | 96 (within-subject) | 1 step: 78<br>½ step: 78<br>¼ step: 77<br>1/8 step: 50 | N/A |
| **PPI assay (Figure 1D/E)** | 100 msec: 72<br>200 msec: 37<br>300 msec: 40<br>400 msec: 34<br>500 msec: 33<br>1000 msec: 38 | 100 msec: 47<br>200 msec: 28<br>300 msec: 34<br>400 msec: 18<br>500 msec: 24<br>1000 msec: 28 | 100 msec: 30<br>200 msec: 20<br>300 msec: 7<br>400 msec: 6<br>500 msec: 6<br>1000 msec: 9 |
| **MK-801 assay (Figure 2A/B)** | DMSO-control: 79<br>MK-801: 84 | DMSO-control: 58<br>MK-801: 72 | DMSO-control: 20<br>MK-801: 51 |

When the above parameters were applied to the prepulse inhibition setting, a clear PPI was observed in larval zebrafish at all ISI values (Figure 1D and E). A one-way between-subjects ANOVA of percentage PPI for each ISI tested was significant (*F*(5, 173) = 3.53, *p* = 0.0046), and post-hoc t-tests revealed that this was driven by a difference in percentage PPI between 200 msec ISI and 300 msec ISI (*p* = 0.0254), 400 msec ISI (*p* = 0.0469) and 500 msec ISI (*p* = 0.0127). Comparing between startle alone and prepulse/startle trials, the PPI effect seems to be caused by the trend for reduced response likelihood in prepulse/startle trials (*M* = 0.27) compared to startle pulse alone trials (*M* = 0.70). Considering startle magnitude, a two-way ANOVA across trial types and ISIs was only significant for trial type (*F*(1, 254) = 19.78, *p*<0.0001), and caused by an increase in response magnitudes between trial types at 300 msec ISI (*p* = 0.0147) and at 1000msec ISI (*p* = 0.0259; data not shown). These findings thus largely concur with previous reports that have measured c-bend startle responses to determine PPI. The PPI percentage reaches a maximum at intermediate ISIs and is driven mostly by reduced response likelihood (not response magnitude) to startle stimuli when they are preceded by a prepulse stimulus.

In order to further validate this assay to be fully applicable to neuropsychiatric models in the zebrafish, we tested whether PPI percentage could be modulated by Dizocilpine (MK-801), a non-competitive NMDA receptor antagonist, commonly used to create animal models of schizophrenia. Larvae were exposed to the drug for the full duration of the PPI assay while controls were exposed to a DMSO-matched control solution. As expected, the PPI percentage was subsequently greatly reduced in MK-801-exposed larvae compared to controls (*t*(128) = 3.563, *p* = 0.0005; Figure 2A). Similarly to the results above, DMSO-control larvae showed a reduced response likelihood between startle alone and prepulse/startle trials (startle alone = 0.73 v. pp/startle = 0.25). However, MK-801 treated fish showed much less difference in response likelihood between trial types (startle alone = 0.86 v. pp/startle = 0.61). In addition, a two-way ANOVA of response magnitudes by trial type and drug condition showed no main effect of trial type (pp/startle v startle alone: *F*(1, 197) = 0.014, *p* = 0.9057) but there was a main effect of drug condition (MK-801 V DMSO-control: *F*(1, 197) = 28.71, *p* = 0.0001), caused by significant elevation in startle response magnitude for the MK-801 group on both types of trials (startle alone *p* = 0.0004; prepulse/startle *p* = 0.0003; Figure 2B). Therefore, MK-801 drove a decrease in PPI percentage through sustaining a high likelihood of response on prepulse/startle trials, though it is also worth mentioning that overall startle magnitude was increased across all trial types in the drug.

**Figure 2.**
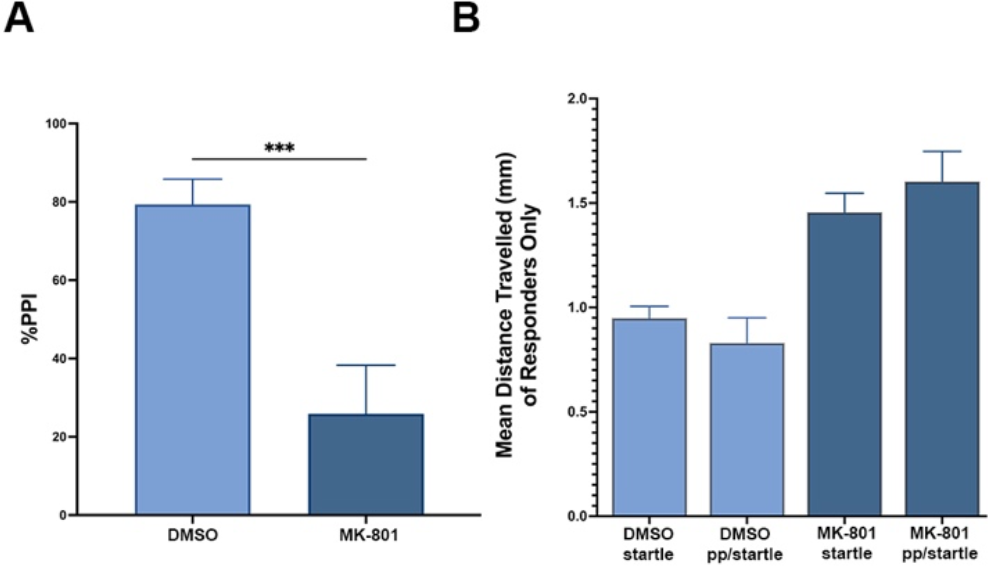
Modulation of PPI response by MK-801. **A**. Startle magnitude of responder larvae in startle alone and prepulse/startle trials for larvae treated with MK-801 or DMSO-controls. **B**. Mean percentage of PPI in responder MK-801 and DMSO-control larvae. For all *N*-numbers, see Table 1 in Methods section. *** = *p*<0.001.

In summary, the data here show that a PPI paradigm can be conducted using larval zebrafish by simply measuring locomotion as a behavioural read-out for startle. Consistent to what has been observed when measuring c-bends in larval zebrafish, our PPI assay reduces the startle likelihood in larval zebrafish whilst the startle magnitude remains the same across prepulse/startle and startle alone trials (Bergeron et al., 2015; Burgess & Granato, 2007).

Our results further showed reductions in PPI by administration of the NMDA receptor antagonist MK-801, in line with previous studies using different PPI startle readouts (Bergeron et al., 2015; Wolman et al., 2011). This is particularly important for creating simpler models, as strong links have been made between glutamatergic abnormality, particularly in the hippocampal CA1/CA3 regions in mammals, and schizophrenia risk/symptoms (Briend et al., 2020; Demjaha et al., 2014; Egerton et al., 2018; Park et al., 2021; Uno & Coyle, 2019). Interestingly, there is a direct translatability in the PPI response between animal models and humans. Notably, the ISI parameters of the PPI assay for larval zebrafish assays are nearly identical to those used with human participants, allowing direct comparability between endophenotypes (San-Martin et al., 2020; Swerdlow et al., 2006).

High temporal resolution PPI that measures larval c-bends are useful for more fine-grained analysis of neuronal circuits in motor movement and therefore might be still selected for certain experimental paradigms (Eaton et al., 1977; Liu & Fetcho, 1999; Medan & Preuss, 2014). However, our simpler locomotion-based PPI assays are clearly suitable to provide a tool for rapid assessment of sensory gating in larvae. This is particularly useful for conducting high-throughput screens of disorder-associated genes and novel compounds, one of the strengths of the zebrafish larval system.

## Materials and Methods

### Zebrafish Husbandry and Maintenance

Zebrafish (Danio rerio) adults and embryos were maintained in accordance with the Animals (Scientific Procedures) Act 1986 under license from the United Kingdom Home Office (PP7266180). AB strain wildtype zebrafish larvae were used at 6 days post-fertilisation (dpf). Embryos and larvae were maintained at 28.5°C on a 14 h ON/10 h OFF light cycle in methylene blue (methylthioninium chloride)/1X Danieau’s medium (NaCl 174mM, KCl 2.1mM, MgSO4 1.2 mM, Ca(NO_3_)_2_) 1.8mM, HEPES 15mM, pH7.6).

### Drug Preparation

MK-801 hydrogen maleate was dissolved in 100% DMSO and then further diluted to 100μM of MK-801 and 0.01% DMSO. The control comparison solution was 0.01% DMSO in Danieau’s medium.

### Prepulse Inhibition (PPI) Assay

All experiments were carried out between 12:00-17:00 in a Zantiks MWP behavioural box (https://zantiks.com/). Larvae were habituated to the procedure room 30 minutes prior to the experiment. All larvae were then placed into a 96-well plate for PPI vibrations and locomotion tracking. Each well was filled with 200μl of Danieau’s medium. In the MK-801 experiment, larvae were placed in 200μl of DMSO-matched control or 100μM MK-801.

Startle stimuli were delivered using a motor mechanism that could produce script-controlled vibrations. The motor was programmed to move in alternating directions, clockwise and anti-clockwise for a total of 4 step movements, each lasting 5 msecs at a 200Hz frequency. This was consistent in all single-pulse and PPI assays shown here. In the process of optimising the PPI assay, the vibration stimuli were presented at varying step sizes (1, ½, ¼ and 1/8^th^ step) where 1 full step size was a 1.8° motor movement. The inter-stimulus interval (ISI) was varied between 100msecs and 1000msecs. In the final PPI assay (see Figure 1C), 8 stimulus trials occurred with a 5-minute inter-trial interval (ITI) and a 300 msec ISI. Trial types alternated randomly between startle stimulus alone and prepulse/startle stimulus trials to avoid habituation to trial types. The startle stimulus was 1 step size and the prepulse stimulus was 1/8^th^ step.

### Data Analysis

Data was extracted using Matlab (R2022B), and statistically analysed and graphed using GraphPad Prism (version 9.5.1). Prior to data analysis, larvae showing 0mm locomotion +/-500 msec around the startle vibration were excluded from further analyses, as was data that was subject to technical errors. PPI percentage was calculated using the following equation:

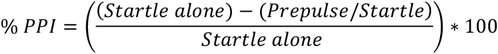

“Startle alone” was the mean locomotion at +100 msecs in startle alone trials and “Prepulse/Startle” is the mean locomotion at +100 msecs in prepulse/startle trials (Figure 1D). 0 msec was the time bin that the startle pulse was delivered in (see Figure 1D). For analyses of % PPI, startle likelihood and startle magnitude, larvae were classified as a “responder larva” if they showed locomotion of >0mm in the +100 msec time bin (i.e., a startle response). *N*-numbers for larvae included in each experiment and analysis type can be found below in Table 1.

## Acknowledgements

We would like to thank Bill Budenberg and Shanelle Kohler (Zantiks) for their advice while optimising the setup of the assay, as well as all the Hindges Lab members for helpful discussions. For the purposes of open access, the author has applied a Creative Commons Attribution (CC BY) licence to any Accepted Author Manuscript version arising from this submission.

## Author Contributions

Emily Read: Investigation, Methodology, Visualisation, Data curation, Formal analysis, Conceptualisation, Writing – original draft, Writing – review & editing.

Robert Hindges: Resources, Methodology, Funding acquisition, Project Administration, Supervision, Writing – review & editing.

